# It is the Locus Coeruleus! Or… is it? : A proposition for analyses and reporting standards for structural and functional magnetic resonance imaging of the noradrenergic Locus Coeruleus

**DOI:** 10.1101/2021.10.01.462807

**Authors:** Yeo-Jin Yi, Falk Lüsebrink, Anne Maaß, Gabriel Ziegler, Renat Yakupov, Michael C. Kreißl, Matthew Betts, Oliver Speck, Emrah Düzel, Dorothea Hämmerer

## Abstract

The noradrenergic locus coeruleus (LC) in the brainstem shows early signs of protein pathologies in neurodegenerative diseases such as Alzheimer’s and Parkinson’s disease. As the LC’s small size (approximately 2.5 mm in width) presents a challenge for molecular imaging, the past decade has seen a steep rise in structural and functional Magnetic Resonance (MR) studies aiming to characterise the LC’s changes in ageing and neurodegeneration. However, given its position in the brainstem and small volume, great care must be taken to yield methodologically reliable MR results as spatial deviations in transformations can greatly reduce the statistical power of the analyses at the group level. Here, we suggest a spatial transformation procedure and a set of quality assessment methods which allow LC researchers to achieve the spatial precision necessary for investigating this small but potentially impactful brain structure.

Using a combination of available toolboxes (SPM12, ANTs, FSL, FreeSurfer), individual structural and functional 3T LC scans are transformed into MNI space via a study-specific anatomical template. Following this, the precision of spatial alignment in individual MNI-transformed images is quantified using in-plane distance measures based on slice-specific centroids of structural LC segmentations and based on landmarks of salient anatomical features in mean functional images, respectively.

Median in-plane distance of all landmarks on the transformed structural as well as functional LC imaging data were below 2 mm, thereby falling below the typical LC width of 2.5 mm suggested by post-mortem data.

With the set of spatial post-processing steps outlined in this paper and available for download, we hope to give readers interested in LC imaging a starting point for a reliable analysis of structural and functional MR data of the LC and to have also taken a first step towards establishing reporting standards of LC imaging data.

## 1. Introduction

The locus coeruleus (LC) is a small nucleus located in the brainstem adjacent to the lateral floor of the fourth ventricle and our major source of noradrenaline in the brain. The noradrenergic LC’s implications for brain function cover a broad range of processes spanning from basic autonomic functions, such as modulation of sleep-wake cycles (Aston-Jones & Bloom, 1981; González & Aston-Jones, 2006); to cognitive functions, such as modulation of attention (Usher et al., 1999; Mather et al., 2016) and memory encoding (Mello-Carpes & Izquierdo, 2013; Sterpenich et al., 2006). Additionally, the LC appears to play a role in neurodegenerative diseases such as Alzheimer’s and Parkinson’s disease, where it is affected early by tau protein pathologies, functional decline, and cell loss (Gesi et al., 2000; Grudzien et al., 2007; Del Tredici & Braak, 2013; Kelly et al., 2017; Betts et al., 2019). Indeed, changes in LC function and structure related to neurodegenerative conditions have been shown in post-mortem studies (Zarrow et al., 2003; Wilson et al., 2013, Theofilas et al., 2017), animal model studies (Arnsten & Goldman-Rakic, 1985; Kalinin et al., 2007), and pharmacological investigations (Rommelfanger et al., 2007). However, precise structural and functional in vivo measurements of the LC in humans are necessary for understanding its relevance as a biomarker in Alzheimer’s and Parkinson’s disease (Betts et al., 2019). Unfortunately, its small size (about 2.5mm in width and 1.5cm in length; Mouton et al., 1994; Fernandes et al., 2012) makes it a difficult target for molecular imaging of tau pathologies which typically operates at a resolution of 3mm^3^. Recent developments in magnetic resonance imaging (MRI) protocols allow us to overcome this limitation and measure LC structure and function *in vivo* with sub-millimetre resolution (Liu et al., 2017; Kelberman et al., 2020 for a review; Betts et al., 2019). Importantly, an MR investigation of the LC comes with the added benefit of allowing for functional assessments of the LC as neuronal function can be assumed to be altered before pathology-related cell loss occurs (Giguère, Nanni, & Trudeau, 2018). However, while *in vivo* structural and functional LC imaging presents as a promising biomarker, acquiring and analysing LC scans is not without its methodological challenges.

The major methodological challenges in functional LC imaging stem from (1) its small size, which necessitates high spatial resolution, high effective contrast in data acquisition, and cautious alignment and spatial normalisation of acquired functional data into the group space, and (2) its position in the dorsal part of the upper brainstem in proximity to the major arteries and pulsatile ventricles, which makes LC imaging prone to physiological noise artefacts from breathing and pulsation (Brooks et al., 2013). Physiological noise and movement correction for imaging small structures has been extensively reported elsewhere. We refer readers to the helpful works of Brooks et al. (2013) and Lawson et al. (2013).

As outlined above, MR imaging sequences with suitable contrasts and resolutions are now available for researchers interested in *in vivo* structural LC imaging in the sub-millimetre range for structural LC imaging (Betts et al., 2017; Priovoulos et al., 2018) and in the 2–0.75 mm range for functional LC imaging (Moeller et al., 2010; Koopmans et al., 2011; Jacobs et al., 2020). However, while current image acquisition tools offer sufficient spatial precision, spatial misalignment in LC applied to the acquired images can still hinder a conclusive interpretation of functional LC imaging results (Liu et al., 2017). For instance, a recent review showed that many reported LC activations spanned far beyond the known anatomical boundaries of the LC (Liu et al., 2017). As illustrated in Figure 1, a group-level analysis that is affected by spatial misalignment can result in reduced statistical power by ‘averaging away’ the LC activations across subjects, if the extent of the misalignment is larger than the approximately 2.5mm width of the LC (Fernandes et al. (2012).

**Figure 1.**
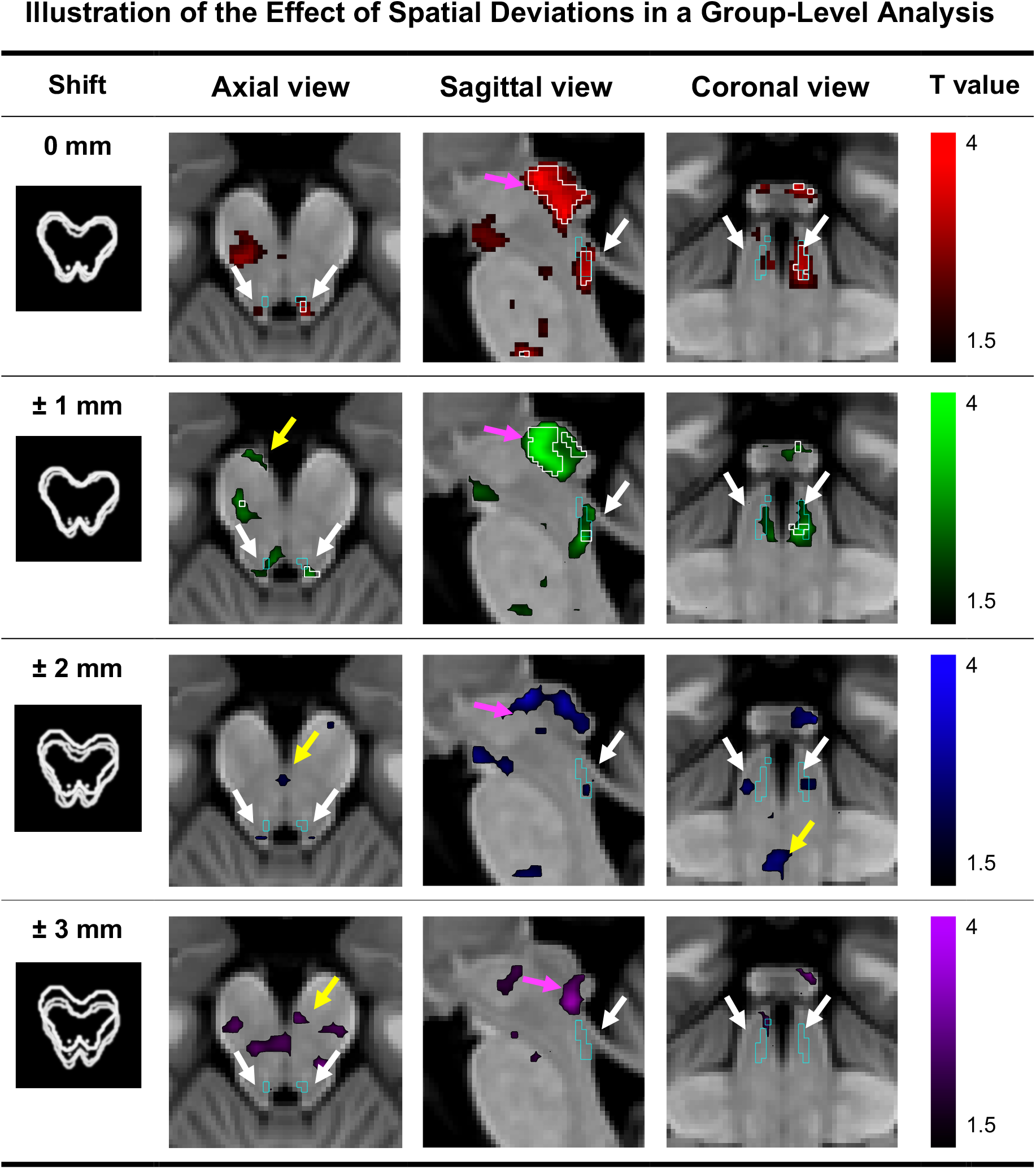
Simulated effects of imprecision in spatial transformations at the group level in the MNI space. Group-level functional activations for reward>non-reward feedback (*p*<0.05, uncorrected; *N*=24, two scans per subject concatenated) are shown using an inclusive brainstem mask. The top row shows the original group-level activation map, while the rows below show the group level result after randomly shifting individual 1^st^ level contrast maps by up to 3mm along the x and y axes to simulate spatial deviations in spatial transformations. White lines indicate significant clusters at voxel threshold *p*<0.005 (the top row [x=-5,y=-38,z=-25; *Z*=2.93; *P*_*FWE-corr*_<0.05, small-volume corrected (SVC) with the meta LC mask by Dahl et al.]; the second row [x=-5,y=-38,z=-24; *Z*=2.73, *P*_*FWE-corr*_=0.062, SVC]; no significant clusters after more than ± 2 mm shifts). As can be seen, activations in the LC area (white arrows) gradually disappear with only minor spatial deviations, whereas spatially more extended activations (pink arrows) are comparatively less affected. Moreover, new spurious activations (yellow arrow) might arise. Cyan-coloured lines show the aggregated meta LC mask created by Dahl et al. (2021).

In this paper, we aim to offer an example of a spatial transformation pipeline based on freely available MR data processing packages which has been adapted to minimise spatial deviations across subjects in the structural and functional LC imaging data for more robust group-level analysis. Additionally, as imaging data quality varies across individuals and studies, we provide a set of quality assessments for spatial processing, which can inform adjustments within the pipeline depending on the dataset and, more importantly, can serve as a reporting standard for processing of LC imaging data. The codes for these analyses and a step-by-step documentation on how to employ them is available for download at https://github.com/alex-yi-writes/LC-SpatialTransformation2021.

The aim of this study was not to compare the precision of different registration and normalisation approaches for functional and structural data as this has been done previously (Ardekani et al., 2005; Klein et al., 2009; Klein et al., 2010; Avants et al., 2011). Several different image processing toolboxes can be used to achieve similar spatial precision for LC imaging (e.g., FSL, SPM, and ANTs). The present study used an intensity-based registration toolbox, ANTs, as the primary method of spatial transformation as *antsRegistrationSyN*.*sh* is known to perform well in multi-modal images as well as non-typical brains such as the atrophied brains of AD patients (Avants et al., 2008).

## 2. Methods and Results

The spatial transformation pipeline and quality assessments are exemplified using a structural and functional imaging dataset adapted for LC imaging (Hämmerer et al., 2018). During functional image acquisition, subjects performed a reward-based memory task. Reward is a highly salient event and was shown to elicit phasic responses of the LC in animal studies (Varazzani, et al, 2015; Glennon et al., 2019).

### 2.1 Participants

Twenty-four healthy younger adults (age range: 20-31 years) were invited to participate in a reward-based memory-encoding task in the scanner. Exclusion criteria included age, past neurological or psychiatric disorders, and the presence of ferromagnetic implants. Each participant was scanned twice as the study compared the effects of two different reward paradigms on memory encoding, resulting in a total of 48 scans. All participants provided written informed consent prior to the experiment and were compensated for their participation and travel expenses.

### 2.2 Task

During the functional scan, participants performed a reward task in which they had to classify the category of a presented picture, for example natural or urban scenery, one of which was associated with a reward, as established in a practice run prior to the scan. A reward or no-reward feedback was presented after the categorization of the picture and was only contingent on the picture type and not on the accuracy of the classification. The two experimental sessions that took place on two different days differed in the proportion of reward and no-reward trials (45 and 135, or 135 and 45, respectively). The association of stimulus category and reward was counterbalanced across participants besides the order of the experimental sessions. Scans were acquired at the German Center for Neurodegenerative Diseases Magdeburg (DZNE Magdeburg).

### 2.3 MRI data acquisition

All images were acquired using a Siemens 3T Biograph mMR scanner (Siemens Healthineers, Erlangen, Germany) with a 24-channel head coil. For each subject, the following images were acquired: a whole-brain T1-weighted MPRAGE anatomical scan to guide functional-structural spatial transformation (1 mm isotropic voxel size, 192 slices, TR = 2,500 ms, TE = 4.37 ms, TI = 1100ms, FOV = 256 mm, flip angle = 7°); a neuromelanin-sensitive T1-weighted multi-echo FLASH sequence for structural LC imaging (0.6×0.6×3 mm voxel size, 48 slices, TR = 22 ms, TA = 4:37, FOV = 230×230×144 mm, flip angle = 23°); and axially oriented T2*-weighted 2D-EPI with Grappa and acceleration factor 2 for functional LC imaging during the reward task (2 mm isotropic voxel size, 51 slices, TR = 3600 ms, TE = 32 ms, FOV = 240×240×102 mm, flip angle = 80°).

### 2.4 Preparing functional and structural MRI data for spatial transformation to MNI space

For each participant, two-session functional images each acquired on different days were first slice-time corrected using the *Slice Timing* function of Statistical Parametric Mapping (SPM12, http://www.fil.ion.ucl.ac.uk/ spm12.html). The output images were then unwarped with distortion fields calculated from the double-echo gradient echo field map and realigned to the mean volume using the *Realign & Unwarp* function of SPM12 in the MATLAB environment using default parameters (Version 2015a, Mathworks, Sherborn, MA, USA, 2015). This generated a mean functional image per person used for spatial transformation of the structural and functional images (cf. Figure 2d). Thereafter, the time-series functional images were smoothed with SPM using a 3×3×3mm kernel in the native space before running single-subject general linear models (GLM) to estimate task-related contrasts in SPM. As physiological noise parameters were not recorded during data collection, they were retroactively corrected using a component-based method (CompCor) during the single-subject GLM analyses (Behzadi et al., 2007). The GLM analysis generated a set of statistical contrast maps in the native space per subject (Figure 2e) that was ready for transformation into the MNI space.

**Figure 2.**
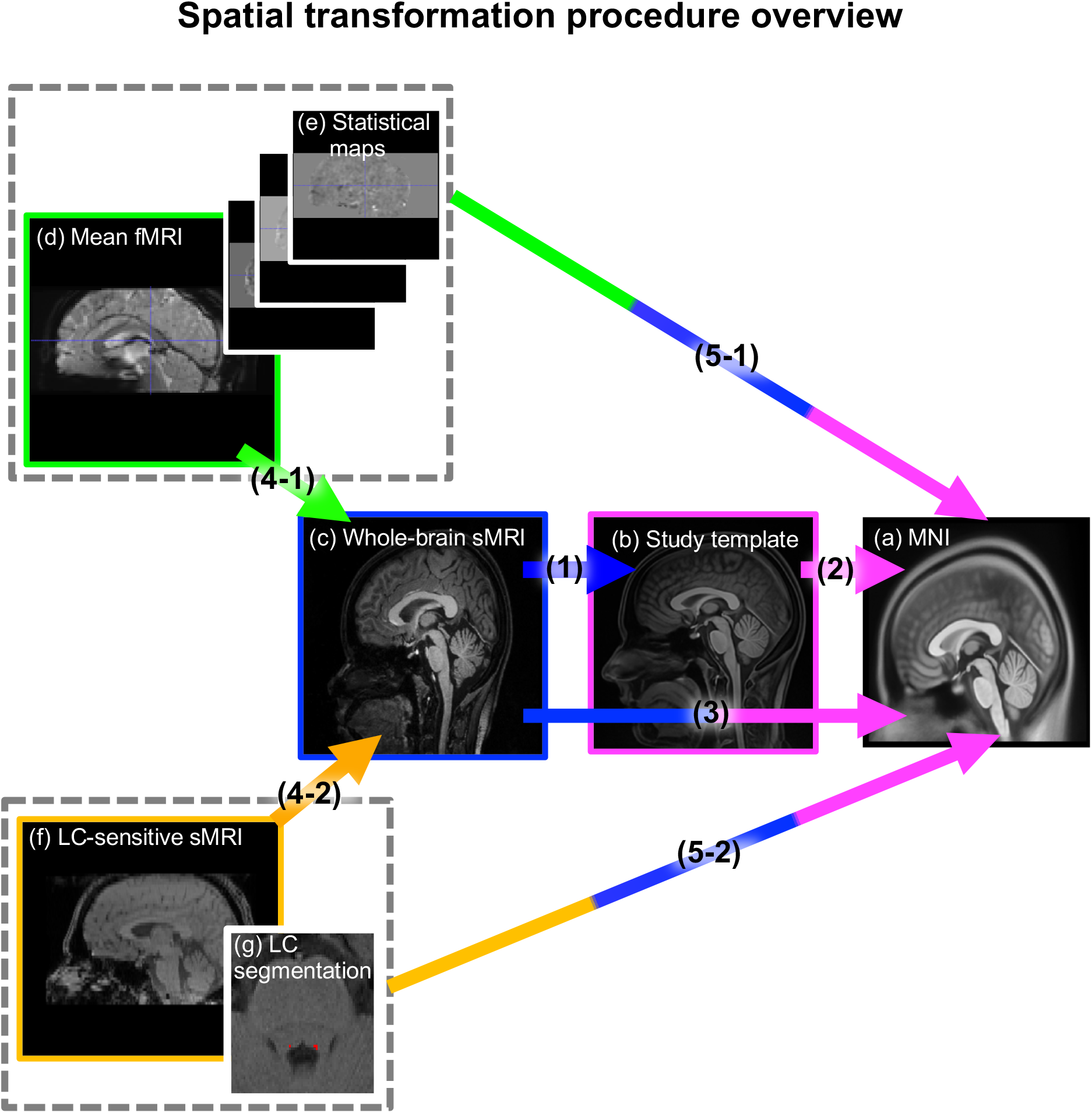
Overview of the spatial transformation steps. Single subject data in native space **(c-g)** are moved into MNI space **(a)** for group-level analyses **(a). (a)** MNI space by Fonov and colleagues (2011). **(b)** A study template space generated from all whole-brain structural images in the dataset was created to help transformation from native to MNI space. **(c)** Whole brain structural image. **(d)** Mean functional image after reslicing, realignment, and unwarping. **(e)** Statistical maps generated from smoothed functional images. **(f)** Neuromelanin-sensitive structural image for LC imaging. **(g)** Manually segmented LC mask drawn on **(f)** in red. Arrows indicate spatial transformation steps, with arrow heads pointing to the image space that an image is transformed into. Triple-coloured arrows indicate concatenated transformation matrices, e.g. the green-blue-magenta arrow (5-1) represents that transformation matrices calculated from a mean functional image (d) to a structural image (c) (step 4-1), structural image (c) to study template (b) and MNI (step 1), and study template (b) to MNI (a) (step 2), are combined in one transformation step (step 5-1). This will transform statistical maps (e) and a mean functional image (d) into MNI space (a) in one transformation step. Numbers indicate the order of transformation steps executed in the pipeline.

Individual T1-weighted whole-brain structural images were bias-corrected using Advanced Normalisation Tools’ *N4BiasFieldCorrection* function (Figure 2c) to correct for RF-field-related inhomogeneity (ANTs, Version 2.3.1; http://picsl.upenn.edu/software/ants/, 2016; Avants et al., 2011; Tustison et al., 2010). A study-specific template (Figure 2b) was created from these bias field-corrected structural whole-brain images using the *antsMultivariateTemplateConstruction2* function of ANTs (Avants et al., 2011).

The LC was manually segmented on the individual neuromelanin-sensitive images (cf. Figure 2f and 2g) using ITK-Snap software (version 3.6.0-RC1; http://www.itksnap.org, 2018) by two independent expert raters (DH and YY). Final LC segmentations contain only the overlapping voxels from both raters (cf. Figure 2g) (Sørensen–Dice coefficient=0.60±0.17; see also Hämmerer et al. 2018 for more details on LC segmentation generation).

### 2.5 Stepwise spatial transformations

Having prepared all relevant data, we outline a procedure which allows for a sufficiently precise spatial transformation of structural and functional LC data to MNI space. As each dataset will vary slightly in contrast properties and signal-to-noise ratios, the parameter settings (i.e., maximal calculations per iteration) of the individual spatial transformation steps might have to be adjusted for specific datasets (see the downloadable manual for details: https://github.com/alex-yi-writes/LC-SpatialTransformation2021).

We used ANTs for all spatial transformations shown in Figure 2 (Klein et al., 2010). However, other suitable spatial transformation tools are available, such as the *Shoot* toolbox from SPM12 (Ashburner & Friston, 2011) or *EPI-reg* function from FSL (Smith et al., 2004).

To facilitate spatial transformation across scans that were acquired at different resolutions, LC-sensitive structural images and LC segmentations (0.6×0.6×3mm native voxel size) were re-sampled to 1-millimetre isotropic voxels matching the MNI space resolution using the *mri_convert* function in FreeSurfer (Version 7.1; http://surfer.nmr.mgh.harvard.edu/, Martinos Center for Biomedical Imaging, Charlestown, Massachusetts). In addition, brain-only binary masks were created from the mean functional images using the *bet* function in FSL (version 6.0.1; https://fsl.fmrib.ox.ac.uk/fsl/, Analysis Group, FMRIB, Oxford, UK) to aid better normalisation by increasing the geometric compatibility among the images.

All spatial transformation steps are shown in Figure 2. First, the individual whole-brain structural MPRAGE image (c) in the native space was registered non-linearly to the study-specific template (b) using *antsRegistrationSyN*.*sh* (step 1). To prepare the transition from the native to the MNI space, the study-specific template (b) was non-linearly registered to the MNI space using *antsRegistrationSyN*.*sh* (step 2 in Figure 2). Individual whole-brain structural images (c) were transformed into the MNI space (a) with the concatenated transformation matrix and deformation fields generated from steps (1) and (2) using *antsApplyTransforms* (step 3). Then, the mean functional image (d) was rigidly registered to the individual structural whole-brain images (c), and the LC-sensitive structural image (f), and the LC segmentation (g) in the space of (f) space were rigidly registered to the individual structural whole-brain image (c) using *antsRegistrationSyN*.*sh* (steps 4-1 and 4-2, respectively). With the concatenated transformation matrices and deformation fields acquired from steps (1), (2), and (4-1), the mean functional images (d) and individual contrast images (e), which are in the same space as individual mean functional images, were transformed to the MNI space (a) non-linearly in one transformation step using *antsApplyTransforms* (step 5-1). The group-level voxel-wise analyses for LC activations can then be performed on the contrast images that have been moved into the MNI space (cf. Figure 1, the first row). Similarly, concatenated transformation matrices and deformation fields acquired from steps (1), (2), and (4-2) were used to transform LC segmentations (g) delineated on neuromelanin-sensitive structural images (f) non-linearly into the MNI space (step 5-2) using *antsApplyTransforms*. All nonlinear spatial transformations were implemented at the 4^th^ degree B-spline interpolation except the individual contrast images (e), which were transformed with the linear interpolation option, and the individual LC segmentations (g), which were transformed with the nearest neighbour option. Note that by following a similar approach but omitting the transformation step to MNI space in the concatenated transformation matrices, group analyses for structural and functional data can also be done in study-specific template space (b) (Suppl. Figure 3). For further details regarding the transformation parameters, see the code downloadable at https://github.com/alex-yi-writes/LC-SpatialTransformation2021.

### 2.6 Quality assessment of the spatial transformations of *functional* LC imaging data

An important step that is omitted in most reports of LC imaging data is the quality control of the spatially normalised images. This step is important even with a rigorous stepwise transformation procedure as outlined above, since variations in data quality might result in spatial deviations in some participants. This may stem from variations in the signal-to-noise ratio or contrast during the acquisition, due to head motion or different subject positioning in the scanner as well as interindividual variability in brain anatomy or signal dropouts. Furthermore, this step will provide a quantification of spatial deviations at the group level which is important additional information when reporting and interpreting group-level results. To check for spatial deviations in individual transformations, a quality control procedure (cf. Figure 3) was carried out on the final outputs (i.e., outputs from steps 5-1 and 5-2 in Figure 2) once all images across subjects were transformed to the MNI space as outlined above.

**Figure 3.**
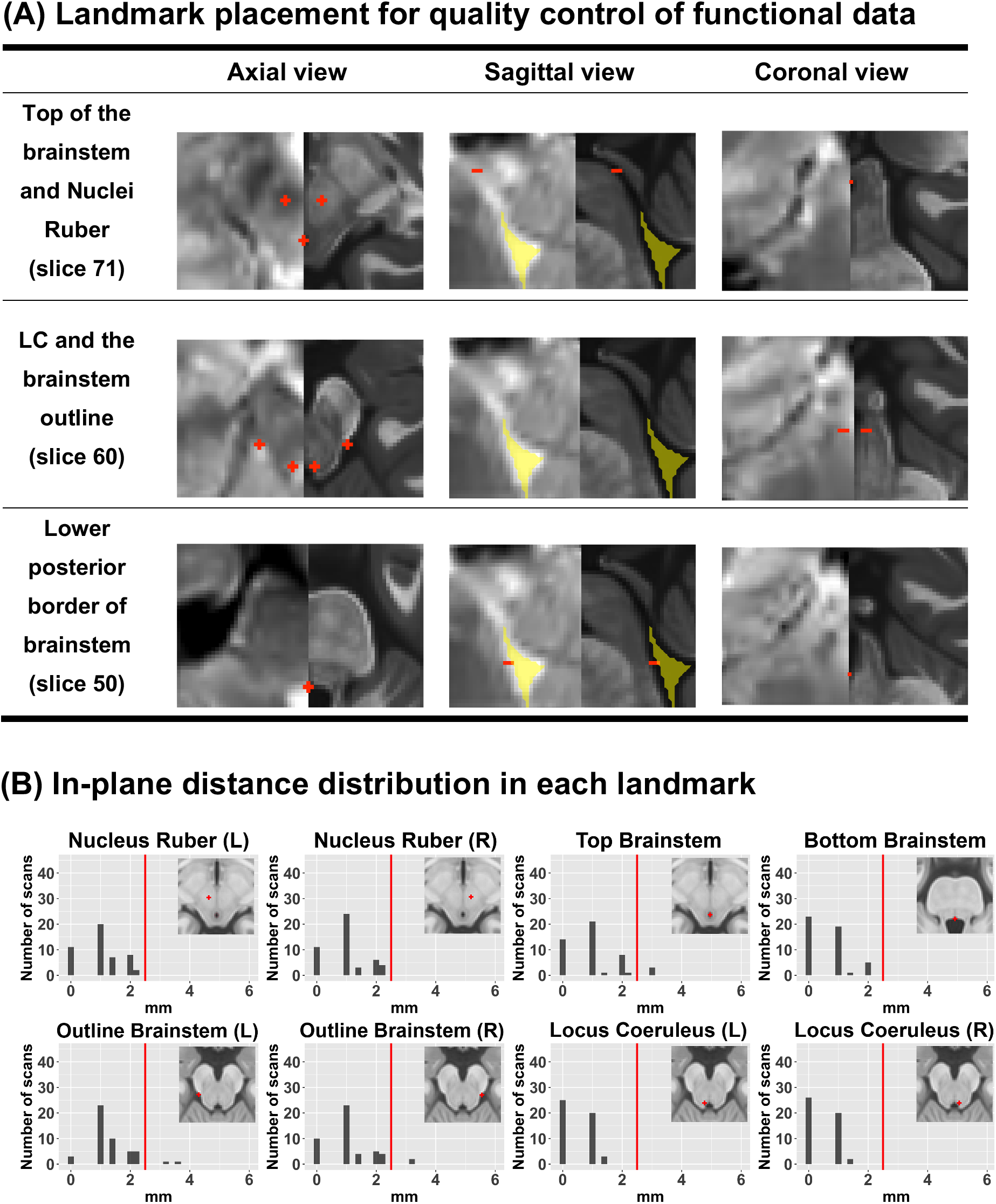
**(A)** Landmarks (indicated by red markers) used in the quality assessment on the individual mean functional images after spatial transformation to the MNI space. Landmarks are drawn on mean functional images (left part of images) before being assessed on the MNI template overlay (right part of images). **(B)** Histograms of in-plane distances between single-subject landmarks and landmarks defined on the MNI template. The median of in-plane distances was at 2mm or lower for all landmarks, thereby falling below the typical width of the LC of 2.5mm (indicated by solid red line, Fernandes et al., 2012). Note that deviations in the outline of the brainstem are bound to differ more from the MNI marks as the precise position along the border of the brainstem is less relevant than capturing the border between brainstem and CSF (cf. row 2 in A). See downloadable manual for details on how to set and evaluate landmarks.

As shown in Figure 3, to assess the transformation precision of the functional images, it is important to establish anatomical landmarks relevant to the LC area in the MNI space, which are evident on the normalised *functional* images (because the LC itself is not visible). Identified landmarks therefore benefitted from anatomical structures that are clearly visible on the mean functional T2*-weighted images, which include the iron-rich nucleus ruber and the border between the cerebrospinal fluid and brainstem of the fourth ventricle (Figure 3, A). By checking the transformation congruence of the mean functional image across subjects, we assessed the transformation precisions of the individual functional images relevant to the statistical analyses (cf. Figure 2, images (d) and (e) are in the same space). Afterwards, spatial transformations are evaluated by overlaying the structural MNI template on the transformed functional image after delineating the landmarks (cf. Figure 3A, right half of pictures). This step also allows for identifying individual cases that might not be well-aligned in the group space, due to interindividual differences in image quality (e.g. low signal intensity in subcortical areas, functional images with high-intensity out-of-tissue areas, or lower signal-to-noise ratio due to suboptimal measurement conditions). These issues can be remedied by additional bias-field correction for functional images or adjusting intensity thresholds of functional mask generation for these individuals (see the downloadable manual for details: https://github.com/alex-yi-writes/LC-SpatialTransformation2021). Landmark checks then must be repeated for each misaligned image after the rectification. It is important to note that landmarks are never to be moved retrospectively once the MNI template has been overlayed.

In the second step, once the transformation quality of all functional images was satisfactory at a single-subject visual inspection stage, the saved landmark images can be aggregated across subjects to assess, quantify and report the quality of spatial transformations at the group-level in MNI space. To quantify the quality of the transformation, landmarks outlined in Figure 3A were also drawn on the MNI space ((a) in Figure 2) in advance to calculate the distance between the MNI-defined landmarks and those that are drawn on the individual transformed mean functional images. The same approach can be applied to the procedure based on the study-specific template analyses. Then, the in-plane distance between each landmark on the individual mean functional images in the MNI space and the landmarks drawn on the MNI space were calculated using a custom MATLAB script downloadable here https://github.com/alex-yi-writes/LC-SpatialTransformation2021. This allowed quantification of the accuracy of the functional image transformation (Figure 3B). As anatomical structures used for landmarks span across several slices, it is recommended to identify the slice number for the respective landmarks on the MNI template before performing quality assessment and using this slice number on the MNI-transformed mean functional images (cf. Figure 3A). To minimise signal loss due to spatial misalignment in the group averages, spatial deviations of the LC landmarks should at least fall below the known width of the LC of 2.5mm (Fernandes et al., 2012).

Figure 3B shows that the in-plane spatial deviations of LC-focused landmarks do not exceed 2.5mm. The observed deviations at this stage of spatial transformation come from various sources such as anatomical idiosyncrasies in each brain or insufficient contrast in the brainstem area. However, they do not seem to originate from the application of nonlinear transformation to the MNI space, as the landmark deviations calculated from the rigid registration step between the mean functional image and structural whole-brain image (step 4-1) shows a similar range of deviations (Suppl. Figure 1).

### 2.7 Quality assessment of the spatial transformations of *structural* LC imaging data

As a first means to assess the spatial precision of the LC segmentations that are moved into the MNI space, aggregated LC segmentations can be plotted as a heatmap, in which a 0 voxel value indicates no shared LC segmentation voxels, while a value of 1 indicates all individual segmentations including a particular voxels. Such a heatmap can also be compared to a meta LC mask in the MNI space as a visual inspection of the coherence and validity of the spatial transformation into the MNI space (cf. Figure 4A).

**Figure 4.**
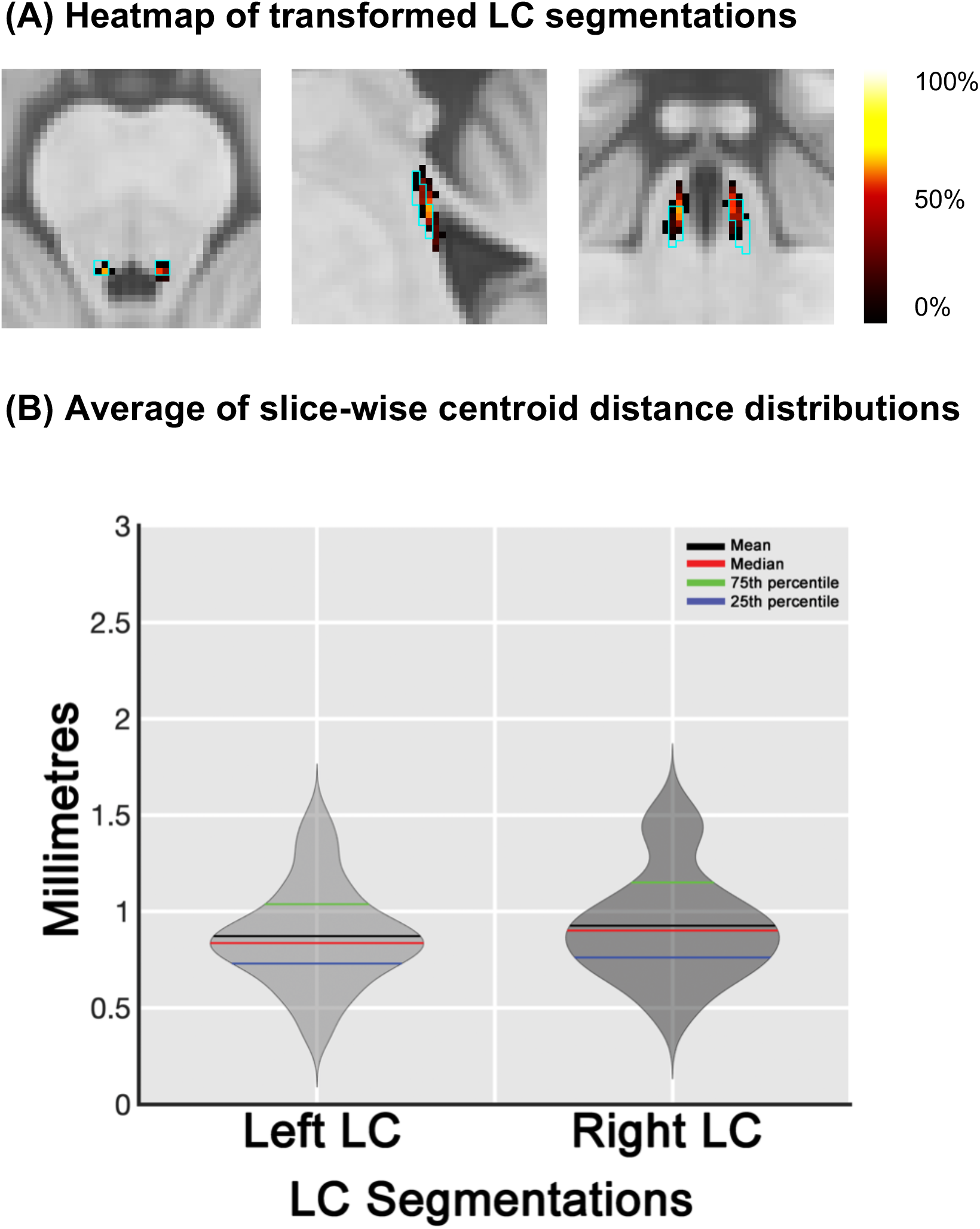
(**A**) A heatmap of transformed individual LC segmentations in the group space. The cyan line indicates the meta LC mask created by Dahl and colleagues (2020). The maximum overlap is at 62.5% and the minimum at 2%. (**B**) Violin plots showing the distribution of distances across subjects for the left and right LC centroid voxels of aggregated meta LC mask and MNI-transformed single-subject LC segmentations. The in-plane distance is calculated slice-by-slice separately for left and right LC and averaged across slices to yield one value per subject and left or right LC segmentation (right: *M*±*SD*=0.92±0.27, *IQR*=0.30; left: *M*±*SD*=0.87±0.25, *IQR*=0.23). A plot showing all cases of slice-wise distances across all LC segmentations and slices can be found in Supplementary Figure 2.

However, the precision of the spatial transformation of the individual LC segmentations should also be quantified by e.g., calculating the slice-wise distance between the centres of each individual MNI-transformed LC segmentation and a template LC mask in the MNI space (e.g., the meta-mask of Dahl et al. (2020)) (cf. Figure 4B). On each slice of the transformed individual LC segmentations, the 3D coordinates of the left and right LC centroids are calculated using a custom MATLAB script downloadable here: https://github.com/alex-yi-writes/LC-SpatialTransformation2021. Likewise, centroid coordinates of the meta LC mask of Dahl and colleagues (2020) were calculated per slice, and distances between these centroid points and the centroid points of the individual segmentations were computed. Then, the distances were averaged across slices within subjects for each side of the LC mask using a custom MATLAB script. The averaged slice-wise distance of the right LC (*N*=48) was 0.70 mm (*SD*=0.21, median=0.64) and that of the left LC (*N*=48) was 0.72 mm (*SD*=0.20, median=0.66), which are both below the width (2mm) of the aggregated meta LC mask created by Dahl et al (2020).

It should be noted that our sample included only younger adults, who are expected to show lower signal strengths in structural LC imaging presumably due to the continuing accumulation of neuromelanin with increasing age (Liu et al., 2019). It might thus be assumed that structural LC recordings in younger adults underestimate the actual volume of the LC as not all LC cells might be sufficiently ‘labelled’ yet. This would explain the pattern of increasing LC signal in neuromelanin-sensitive scans during adulthood (Liu et al., 2019). Furthermore, as our focus in this paper was to provide a precise localisation of the LC, the LC was segmented using conservative thresholds, which thereby most likely underestimated the actual LC volume. Finally, our analyses (Suppl. Figure 2) and postmortem data (Fernandes et al., 2012) show that also individual LC positions in their native space vary in the range of 2.7–6.7mm (*M*±*SD*=4.3±1.0mm) with respect to the posterior midline of the brainstem across individuals due to interindividual differences in anatomy (cf. Figure 5, orange plots on the left side of the figure).

**Figure 5.**
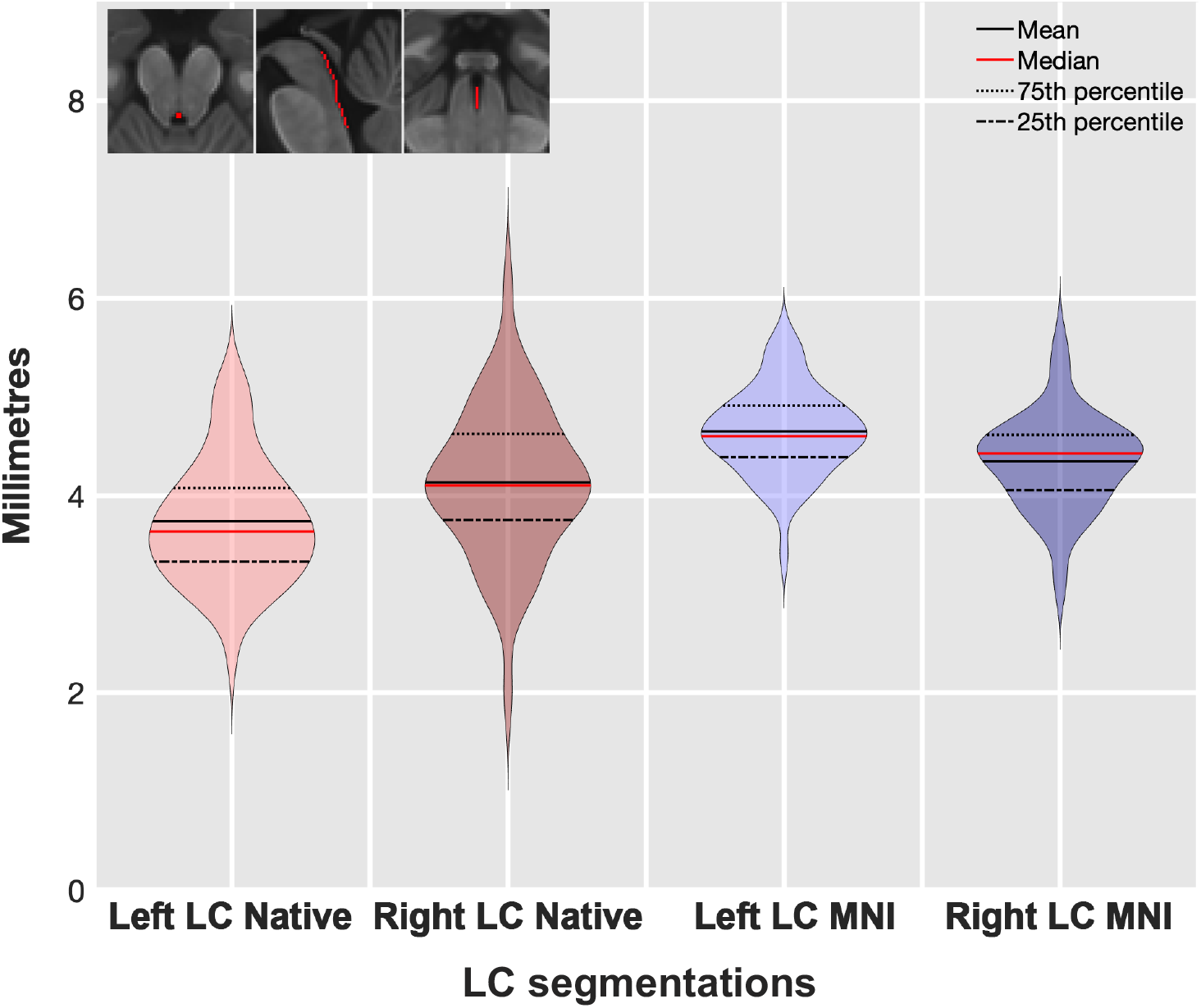
Mean slice-wise Distance of centres of individual LC segmentations with respect to brainstem midline in native and MNI space. Each violin plot indicates slice-wise distance from either left or right LC segmentation centroid coordinates (native or transformed) to the midline coordinates. Midline coordinates have been manually defined on the posterior border of brainstem and the 4th ventricle (c.f. inset). Distances are assessed *and* averaged across slices per subject. Mean distance of LC from brainstem midline (and *variability in* this distance) across individuals are as follows from left to right: Left LC segmentation resampled at 1mm isotropic voxel in the native space (M±SD=3.74±0.61, median=3.64), and right LC (M±SD=4.13±0.75, median=4.10), Left MNI-transformed LC segmentation (M±SD=4.65±0.43, median=4.60), right MNI-transformed LC segmentation (M±SD=4.35±0.48, median=4.43).

This is corroborated by post-mortem studies that also report deviations in individual LC positions and size (German et al., 1988; Fernandes et al., 2012). As normalisation to the group space is based on discernible features of the brain anatomy such as ventricles and outline of the brainstem, variations in LC position between individuals in the native space will to some extent translate to LC segmentations in the group space (violet plots on the right side of Figure 5). Therefore, the delineation of the LC combined across all subjects might exceed the typical size of an individual LC and be prone to varying levels of overlap across indinviduals. This current obstacle of assessing LC segmentations at the group level will greatly benefit from probabilistic LC masks (Ye et al., 2021), which eventually are expected to incorporate age- and disease-related information on partial volume and neuronal loss effects in specific areas of the LC in certain populations.

#### Box 1.

**Suggestions for reporting registration and normalisation precision for functional LC imaging**

Following our outline of an analysis pipeline and a set of quality checks on the precision of spatial transformations for functional and structural LC imaging, we propose the following standards for reporting group-level LC imaging results. Many existing publications already include information on some of the aspects outlined below. However, very few encompass all aspects outlined here, especially when it comes to including information on quality assessments of spatial transformations. As shown in Figure 1, our ability to reliably identify LC activations at the group level is crucially dependent on the precision of the post-hoc spatial transformations of the functional images. Thus, we would like to propose the following information to be included as a reporting standard of LC imaging studies:

1. *In-plane distances of landmarks drawn on each subject’s mean functional images in the MNI space from pre-defined landmarks on the structural MNI or group template image* (cf. Figure 3).
2. *Slice-wise distances between the centre of an LC template mask and the centre of each subject’s LC mask in MNI or group template space* (cf. Figure 4). Moreover, given the LCs’ position in the brainstem and its small size, the following information on data preprocessing should be given:
3. The description of the *movement correction method* should include replicable details and should mention any deviations from default settings or additional correctional techniques performed.
4. The description of the *physiological noise correction method* should include replicable details and mention any deviations from default settings or additional correctional approaches taken. If no physiological parameters have been recorded, independent component analysis (ICA) approaches can be used to achieve similar effects (Beckmann & Smith, 2004).

## 3. Discussion and conclusion

Over the last decade, there have been substantial advances in functional and structural LC imaging, both with respect to novel imaging protocols (Keren et al., 2009; Betts et al., 2017; Priovoulos et al., 2018; Trujillo et al., 2019; Jacobs et al., 2020) as well as in with respect to gaining a better understanding of the contribution of the LC to cognition, behaviour, and neurodegenerative diseases (Betts et al., 2019; Poe et al., 2020; Kelberman et al., 2020). In this article, we suggest a set of spatial transformation and quality assessment steps that can serve as an analysis and reporting standard to further support these advances in the field of LC imaging.

With the set of procedures outlined here, we aim to provide readers interested in LC imaging a mostly plug-and-play approach for processing group-level functional and structural LC imaging data. In this study, we used ANTs, SPM12, FSL, and FreeSurfer for pre-processing and spatial transformations. However, the general aspects of the analysis steps outlined here (cf. Figure 2–4) can be applied to other toolboxes such that other combinations of toolboxes can be employed depending on the user’s preference and proficiency. These include the following: (1) the measures that can be performed for preparing images for the spatial transformation, such as bias correction of images before spatial transformations to compensate for low imaging contrast in the brainstem region, (2) the order of spatial transformations outlined in Figure 2, and (3) using different image registration metrics and varying their parameters (e.g., cross-correlation, mutual information) to optimise registration in individual problematic data points.

Most importantly, we hope that the quality assessments and reporting details of spatial transformations for functional and structural LC imaging data outlined in this paper can serve as a starting point for a reporting standard of LC imaging data.

The focus of this paper is on allowing sufficiently congruent spatial alignment for interpreting group-level results in functional LC imaging. It should be noted that a variety of factors can contribute to spatial deviations at the group level, and not all the deviations can be reduced by improving the spatial alignment. In structural LC imaging, which currently provides our best spatial resolution for LC imaging (Betts et al., 2017; Priovoulos et al., 2018), existing imaging sequences most likely do not capture the entire extent of the LC, which can lead to inaccurate delineation of LC. This can be due to movements during image acquisition which obscures hyperintense LC voxels, partial-volume effects, and insufficient signal for neuromelanin-sensitive imaging in the younger age group (the LC contrast intensity shows an inverted-U shape curve across the lifespan, which may be due to neuromelanin increase during maturation and cell loss during ageing, Manaye et al., 1995; Liu et al., 2019), as well as differences in the properties of the imaging sequences (Betts et al., 2019). Furthermore, the LC position in the brainstem varies across individuals (German et al., 1988; Fernandes et al., 2012), which can translate into spatial deviations at the group level (Figure 5). In the future, examining such group differences can be further assisted by the development of probabilistic LC atlases (Ye et al., 2020), which could help identify and interpret partial LC loss in MNI-transformed individual LC segmentations as well as base transformations on the informed LC shapes of different populations.

Regarding functional LC imaging, limitations in the intersubject image alignment of spatial transformations are also related to the inevitably larger voxel sizes (with a currently minimum size of 1.5–2 mm in 3T scanners and 0.75–1 mm in 7T scanners) and smoothing kernels, as well as the lower level of anatomical information available in mean functional images as compared to that of structural data for assessing the spatial precision based on landmarks. Based on these constraints, spatial deviation of at least up to a voxel size in spatial transformation of functional data can often be expected (cf. Figure 3). Given the detrimental effects of spatial deviations on functional LC activation at the group level (cf. Figure 1), functional LC sequences that aim for lower voxel sizes may help to counteract this effect. Nonetheless, as exemplified in Figure 1, also LC imaging studies using larger voxel sizes of up to 3 mm can benefit greatly from improved spatial precision in image processing.

In addition, our sample was rather unconventional as the same subjects were invited twice for MRI scans, which might have resulted in lower noise levels than studies which did not perform repeated assessments. However, this does not affect the principles of the analyses and quality assessments presented in this study. Similarly, although ICA-based methods of correcting for physiological noise have been shown to be comparable in their ability to correct for noise as compared to regression-based approaches using recorded physiological parameters (Salimi-Khorshidi et al., 2014; Griffanti et al., 2014), physiological noise correction using additional recordings for physiological noise is generally preferable. The spatial precision in alignment achieved with our dataset might thus be further improved in a dataset using concurrent acquisition of physiological signal and MR data or higher spatial resolutions in MR data acquisition. However, it might be important to also show that sufficient spatial precision can be obtained in a non-optimal dataset as the present given that not all set-ups might allow for e.g., additional physiological signal recordings. Finally, identifying reliable LC activation depends not only on the precision of spatial transformation, but also on the sample size and the effectiveness of experimental manipulations. However, it is only by starting with methods that allow for a robust assessment of LC activations, as outlined here, that one can address questions regarding which sample sizes or paradigms are most suited for LC imaging.

Brainstem imaging has only recently begun to attract attention for methodological scrutiny (Sclocco et al., 2018; Brooks et al., 2013). Nevertheless, it will in the future undoubtedly afford higher spatial precision than that reported in this paper, owing to new imaging sequences and further developments in spatial transformation methods. We hope that the set of analyses, quality assessments, and reporting standards outlined in this paper can contribute to this development. Finally, although we have been focusing on optimising spatial transformations for brainstem imaging, the analysis steps and quality checks outlined here can be easily adjusted for other small brain areas, such as hippocampal subfields or other small nuclei within or outside the brainstem, for example, habenula or mamillary bodies.

## Acknowledgements

This research was supported by Sonderforschungsbereich 779, Project A07, Sonderforschungsbereich 1315, Project B06, Sonderforschungsbereich 1436, Project A08. DH is funded by ARUK SRF2018B-004.

## Authour contributions

YY, FL, OS, DH, and ED contributed to the conceptualisation and methodology of the study. YY contributed to investigation, analysis, visualisation, and writing of the original draft. DH and ED supervised the investigation, analysis, and writing of the original draft. All authours reviewed and edited the manuscript.

