## Supplementary materials for "It is the Locus Coeruleus! Or… is it? : A proposition for analyses and reporting standards for structural and functional magnetic resonance imaging of the noradrenergic Locus Coeruleus"

### Supplementary Results

1. **The functional-structural image registration quality assessment**

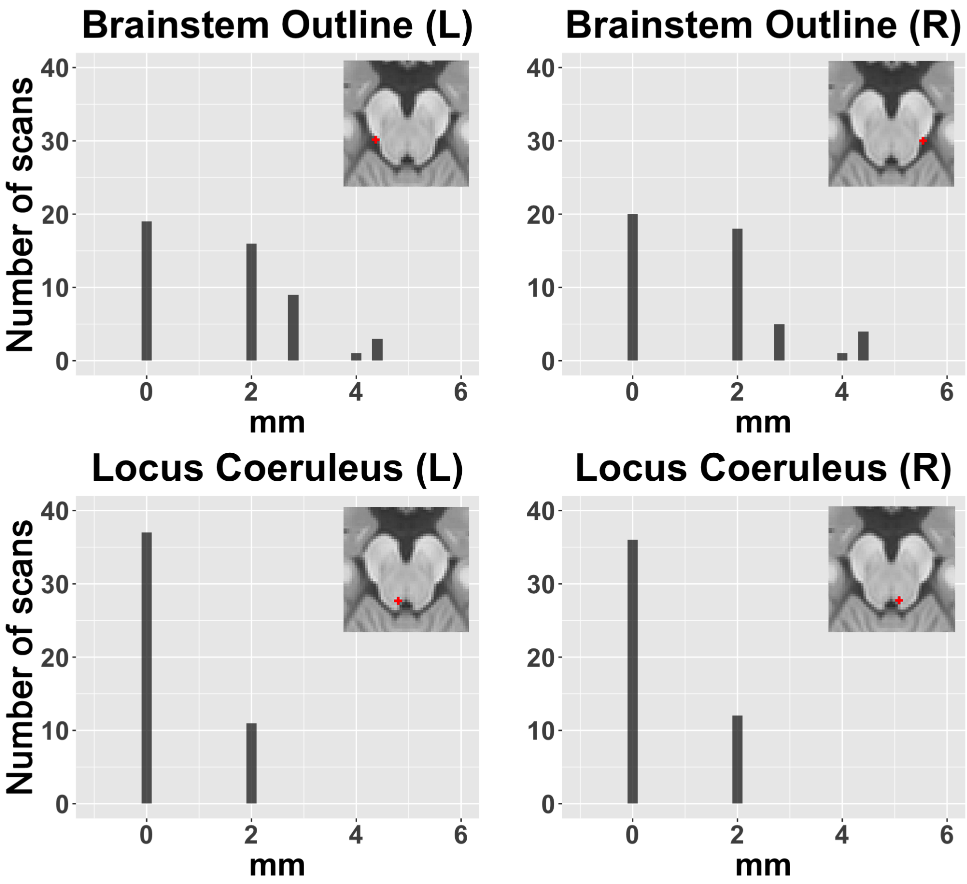


**Supplementary Figure 1**. **In-plane distance of LC-focused landmarks after registration of the native mean functional images and the whole-brain structural images (step 4-1, Figure 2 of the main text).** The in-plane distance was computed by calculating the distance between the manually drawn landmarks of the mean functional images in their native space and the corresponding landmarks in the whole-brain structural images in the mean functional image space in each subject. Note that the distances of the outline of the brainstem between the functional and structural images differ more from that calculated in the images transformed onto the MNI space, as the position along the border of the brainstem varies greatly in the native space (c.f. Figure 3A, the second row of the main text).

The above figure summarizes transformation precision for the registration of individual structural and functional data (c.f. step 4-1, Figure 2 of the main text) prior to normalising. Spatial transformation inaccuracies at this step come from various sources as described in the main text. The above precision assessment of this registration step shows that the nonlinear transformation to MNI space does not add significantly to the spatial deviation observed in the normalised data (c.f. Figure 3B of the main text).

### 2. Distribution of slice-wise centroid distance in all slices in all segmentations

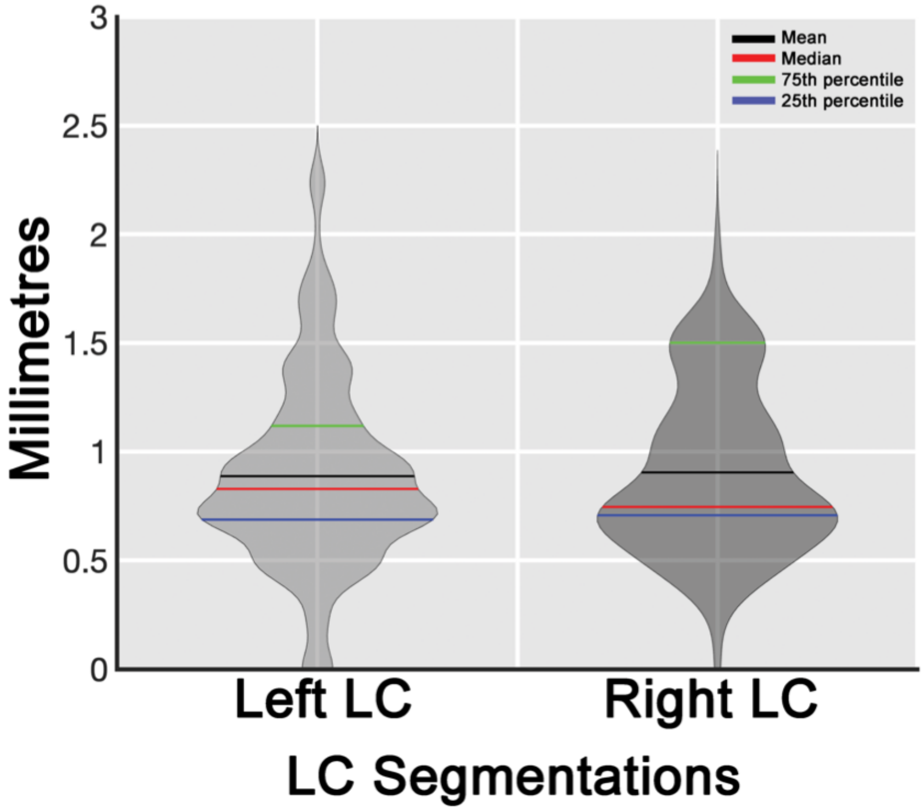


**Supplementary Figure 2**. **Slice-wise distance of all LC segmentation centroids in MNI space (step 5-2, Figure 2 of the main text).** Violin plots showing the distributions of the slice-wise distance in each centroid in each individual segmentation slice (right: *M*±*SD*=0.90±0.37, *IQR*=0.43; left: *M*±*SD*=0.88±0.43, *IQR*=0.41). The method for calculating the distance is described in the main text (Figure 4).

3. **In-plane distance distribution in each landmark in mean functional images transformed onto the study-specific template**

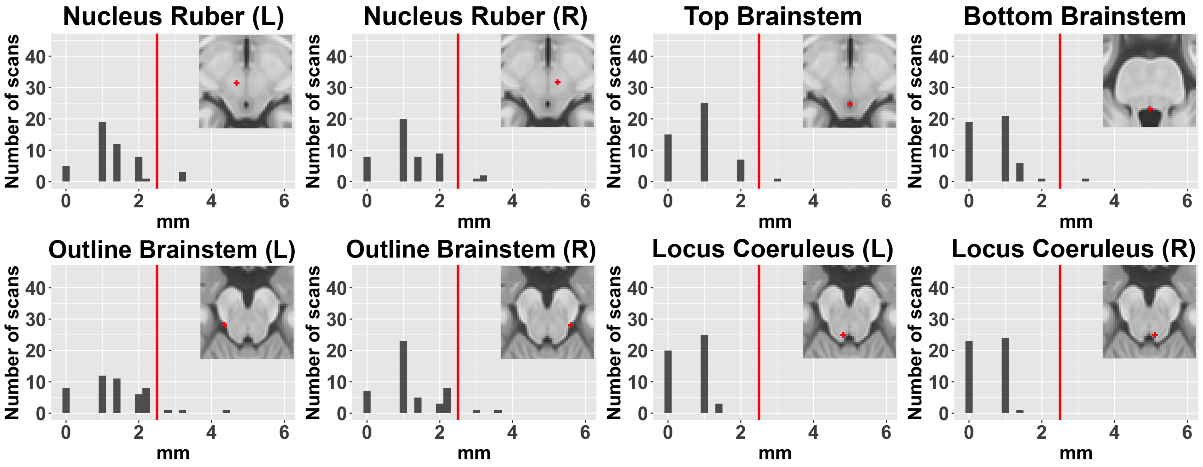


**Supplementary Figure 3. Histograms of in-plane distances between single-subject landmarks and landmarks defined on the study-specific template.** In the study-template-transformed mean functional images, median of in-plane distances of all landmarks were below 2.5 mm, thereby falling below the typical width of the LC (indicated by solid red line, Fernandes et al., 2012). Two paired-samples t-tests were conducted, and, in both LC-focused landmarks (bottom row, middle right and far right), the distances calculated from the study-template-transformed mean functional images and the MNI-transformed images did not show statistically significant difference (**left LC**: [study-template landmark distances: *M*±*SD*=0.61±0.53], [MNI landmark distances: *M*±*SD*=0.55±0.61] *t*(47)=-0.52, *p*=0.60; **right LC**: [study-template landmark distances: *M*±*SD*=0.53±0.51], [MNI landmark distances: *M*±*SD*=0.52±0.60] *t*(47)=-0.11, *p*=0.92).

Depending on the analysis objectives, the spatial transformation procedure and quality assessment of the transformed images can be performed in the study-specific template space.

### 4. A brief visual screening of structural image spatial transformation quality across subjects

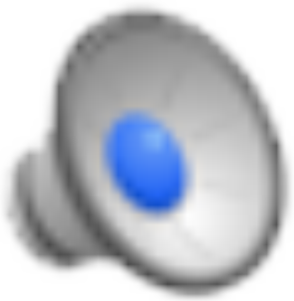

The video is also downloadable here: https://github.com/alex-yi-writes/LC-SpatialTransformation2021/blob/91fd7935a6024a89d6b987940ded4433b73eb787/T1_registryCheck.avi

The video shows the MNI-transformed individual whole-brain structural images (step 3 of Figure 2 of the main text) displayed one after the other to compare their position in MNI space. The slight ‘movements’ on the back of the brainstem (see the red line for a stationary reference) across individual structural images indicates the precision of spatial transformations. Creating this video can be an easy method for spotting individuals with imprecise spatial transformations. The code for creating the video can be downloaded from this link: https://github.com/alex-yi-writes/LC-SpatialTransformation2021
